# In silico DNA codon optimization of the variant antigen-encoding genes of diverse strains of Nipah virus

**DOI:** 10.1101/2021.04.23.441071

**Authors:** Aeshna Gupta, Disha Gangotia, Kavita Vasdev, Indra Mani

## Abstract

Due to an increase in human and wildlife interaction, more and more zoonotic diseases are emerging. A prime example of this is the emergence of the Nipah virus (NiV). Due to high rate of mortality specifically in India and Bangladesh, there is an urgent need for accelerated research for NiV involving the development of vaccines or drugs. The genome of NiV consists of six genes (N, P, M, F, G and L) encoding yielding nucleoprotein, phosphoprotein, matrix, fusion, glycoprotein and large RNA polymerase. We have used these six genes for in silico assessment of DNA codon optimization in *Escherichia coli*. It was observed that the codon adaptation index (CAI) and GC content of the genes in optimized DNA were enhanced significantly as compared to wild-type strain. On an average, CAI and GC content of N gene in optimized DNA was enhanced by 2.3 (135.1%) and 1.2(9.9 %) fold respectively, while in P/V/C it was increased by 2.0 (98.3 %) and 1.1(7.8%) fold respectively. Further, the CAI and GC content in optimized DNA of M gene and F gene was enhanced by 2.0(99.0%) and 1.1(7.2%) fold respectively for gene M and 2.4(142.5 %), 1.2(15.4%) fold respectively for gene F. Gene G showed an increase of 2.1(114.8 %) fold for CAI, 1.1(11.2%) fold for GC content and gene L showed an increase of 2.4(143.7%) fold for CAI, 1.2(17.2%) fold for GC content. Our result demonstrates that the optimized genes could be useful for better expression in host without any truncated proteins and also useful for protein folding and function.

## Introduction

Emerging zoonotic diseases are the products of socioeconomic and anthropogenic environmental changes, Nipah virus (NiV) being one of its best examples. Nipah is a zoonotic disease caused by Nipah virus. Fruit eating species such as Pteropus bats, popularly known as flying foxes, are supposed to be the natural hosts of the virus. NiV emerged as a new virus, causing severe morbidity and mortality in both humans and animals exactly 20 years ago and destroyed the pig-farming industry in Malaysia, and it continues to cause outbreaks in Bangladesh and India [1]. NiV is the second member of the genus *Henipavirus* in the family *Paramyxoviridae* [1]. Similar to other paramyxoviruses, NiV particles are pleomorphic, spherical to filamentous, having an RNA genome and range in size from 40 to 1,900 nm. Among the NiVs known to cause disease in humans, there are two major genetic lineages, i.e., NiV Malaysia (NiV-MY) and NiV Bangladesh (NiV-BD). Genome of the Malaysia NiV is 18,246 nucleotides (nt) in length, whereas that of the Bangladesh NiV is 18,252 nt [2].

The core of the virion contains a linear ribonucleoprotein (RNP) comprising of negative sense single stranded RNA. The genome consists of six genes (N, P, M, F, G and L) encoding nucleoprotein, phosphoprotein, matrix, fusion, glycoprotein and large RNA polymerase [3]. Nucelocapsid protein (N) is the most abundant protein present and necessary for capsid structure. Phosphoproteins (P) and large polymerase proteins (L) aid RNA polymerase in transcribing RNA to mRNA to antigenomic RNA. Traditional lipid bilayer envelopes the virion but it is “spiked” with fusion (F) and receptor-binding glycoproteins (G). Matrix proteins (M) are present on the underside of the lipid bilayer for structural support and regulating the budding process. The P gene encodes at least three nonstructural proteins (C, V, and W) in addition to the P protein. However, P protein is the only essential gene product for genome replication [3]. NiV entry and cell-to-cell spread are driven by two transmembrane glycoproteins, the attachment (G) and the fusion (F) proteins, that are exposed on the surface of viral particles and on infected cells to mediate attachment to the host cell receptor and membrane fusion, respectively [4].

As the world continues to struggle with the COVID-19 pandemic caused by SARS-CoV-2, it becomes all the more crucial to study the characteristics of Nipah virus that might increase its risk of causing a global pandemic in future. The route of infection of NiV from bats to humans is by ingestion and consumption of NiV-contaminated or partially eaten fruits, or by contact with infected animals such as pigs, cattle and goats [5]. Clinical presentation ranges from asymptomatic infection to fatal encephalitis [1]. As an RNA virus, it has an exceptionally high rate of mutation and if a human-adapted strain were to infect communities in South Asia, high population densities and global interconnectedness would rapidly spread the infection [6]. Hence there is a potential need for development of highly immunogenic vaccine against the virus.

Currently, no drugs or vaccines exist for this virus, though many trials are in progress [7]. DNA vaccines also known as ‘naked DNA’ or nucleic acid vaccine, encode antigens of pathogenic organisms including viruses, bacteria, fungi and parasites [8]. However, these are shown to have low immunogenic properties in larger species such as primates and humans [9]. One promising approach for enhancing its immunogenicity is to maximize its expression in the immunized host [10]. Many organisms including viruses tend to have biases towards certain synonymous codon and codon pairs in their genes. A gene containing rarely used codons in one particular organism will show increased expression levels in heterologous system through codon optimization. There are many software tools and technologies which have been developed for gene expression studies and predicting the expression level of genes through computational methods. This is appealing as expensive and difficult experiments are not required [11].

DNA codon optimization is one such useful technology for improving the yields of expressed heterologous proteins. It is a technique to exploit the protein expression in living organism by increasing the translational efficiency of gene of interest by transforming DNA sequence of nucleotides of one species into DNA sequence of nucleotides of another species like plant sequence to human sequence, human sequence to bacteria or yeast sequences [12]. Variation in codon usage is considered as one of the important factors affecting protein expression levels, since the presence of rare codons can reduce the translation rate and induce translation errors with a remarkable impact on the economics of recombinant microbe-based production processes [13,14,15]. Methods for optimizing genes are sophisticated and becoming increasingly popular for a variety of applications such as expression in prokaryotes, yeast, plants and mammalian cells [16]. The host specific epitopes have been earlier identified in influenza A virus [12]. Codon optimization of the Ag85B gene which encodes the secretory antigen of *Mycobacterium tuberculosis* has also proved beneficial [10]. Furthermore, the use of codon optimized genes has allowed notable increases in the production of many enzymes in a variety of hosts, including cellulases in *Saccharomyces cerevisiae* [17], phytases in *Aspergillus oryzae* [18], cutinases [19], lignocellulases [20], and lipases [21] in *Pichia pastoris* and calf prochymosin in *Escerichia coli* [22]. Thus, it can be concluded that the list of products obtained by the expression of codon optimized genes in microorganisms is constantly growing and includes biofuels, pharmaceuticals, novel bio-based materials and chemicals, industrial enzymes, amino acids, and other metabolites [13].

Therefore, the aim of this study is to optimize the codons for over expression of all six target genes of NiV in *E. coli* using in silico tools for production of adequate amount of protein. The synonymous codons were specifically altered without any changes in the amino acid sequence so that antigenicity and functional activity of each protein remains exactly similar to its native type. The DNA codon optimization of the studied genes will be useful in increasing the expression level of desired proteins so as to ensure their efficient production for immunotherapy and immunodiagnostics purposes, without any bias.

## Methods

### Collection of sequences

Nucleotide sequences (cds) of different genes of Nipah virus (Accession number: NC_002728.1, FJ513078.1, AY988601.1, AJ627196.1, AY029768.1, and MH523642.1) were retrieved from NCBI-GenBank (http://www.ncbi.nlm.nih.gov).

### Codon optimization and analysis

Optimizer (http://genomes.urv.es/OPTIMIZER/) [23] is an on-line PHP application useful for predicting and optimizing the level of expression of a gene in heterologous gene expression host. It was used for optimization and calculation of codon adaptation index (CAI), G+C and A+T content of the retrieved DNA sequences with reference to *E. coli*K-12 MG1655 as it is a popular host for heterologous gene expression. CAI was also calculated for each gene of six different strains.

### Statistical analysis

GraphPad Prism (version 8.1) software was used for statistical analysis of genes to calculate mean, range and standard deviation. The values were tabulated and a graph was then plotted to compare the CAI of wild-type and optimized gene sequences among different strains of Nipah virus. The CAI, GC and AT of all 6 genes of Nipah virus were compared using Mann Whitney test. A two-tailed probability p < 0.05 were considered to be statistically significant.

### Nucleotide sequence alignment

Nucleotide sequence alignment was carried out using ClustalW between wild-type and optimized sequences for all 6 genes of strain NC_002728.1 FJ513078.1, AY988601.1, AJ627196.1, AY029768.1, and MH523642.1.

## Results

Currently, no drugs or vaccines exist for Nipah virus, though many trials are in progress [7]. Treatment is limited to supportive care [24]. Ribavirin has shown some evidence for a reduction in mortality, but its efficacy against NiV disease has not yet been established [25]. Therefore, there is an urgent need for development of effective vaccines for which the current study, was undertaken using the DNA codon optimization method for producing adequate quantity of protein in the desired host. The codon usage for various genes of Nipah virus i.e. nucleocapsid protein, P/V/C, matrix protein, fusion protein, attachment glycoprotein and polymerase were summarized in **Table 1-6** respectively. Their codons were optimized with reference to *E. coli*.

**TABLE 1:** Expression level of N (Nucleocapsid) gene of Nipah Virus in *E. coli* of wild-type and codon-optimized sequences.

| NIPAH VIRUS STRAINS (GenBank Accession No.) | WILD-TYPE DNA |  |  | OPTIMIZED DNA |  |  |
| --- | --- | --- | --- | --- | --- | --- |
|  | CAI | GC % | AT% | CAI | GC % | AT% |
| NC_002728.1 | 0.242 | 45.5 | 54.5 | 0.569 | 50.0 | 50.0 |
| FJ513078.1 | 0.244 | 45.4 | 54.6 | 0.571 | 49.8 | 50.2 |
| AY988601.1 | 0.243 | 45.2 | 54.8 | 0.568 | 49.6 | 50.4 |
| AJ627196.1 | 0.237 | 45.5 | 54.5 | 0.568 | 50.2 | 49.8 |
| AY029768.1 | 0.242 | 45.5 | 54.5 | 0.569 | 50.0 | 50.0 |
| MH523642.1 | 0.242 | 45.5 | 54.5 | 0.567 | 49.8 | 50.2 |
| N | 6 | 6 | 6 | 6 | 6 | 6 |
| Min. | 0.237 | 45.2 | 54.5 | 0.567 | 49.6 | 49.8 |
| Max. | 0.244 | 45.5 | 54.8 | 0.571 | 50.2 | 50.4 |
| Mean±SD | 0.242<br>±0.002 | 45.4<br>±<br>0.121 | 54.6<br>±<br>0.121 | 0.569<br>±<br>0.001** | 49.9<br>±<br>0.210** | 50.1<br>±<br>0.210** |

\*\*– p <0.002: in comparison with wild type.

In the present study, we observed that the CAI of optimized sequences was more in comparison to the wild-type sequences. The CAI, GC and AT frequencies in six strains of wild-type nucleocapsid protein ranged from 0.237 to 0.244, 45.2 to 45.5 and 54.5 to 54.8 respectively with an average (±SD) of 0.242(±0.002), 45.4(±0.121) and 54.6(±0.121) respectively. The respective frequencies of these in optimized DNA range from 0.567 to 0.571, 49.6 to 50.2 and 49.8 to 50.4 with an average (±SD) of 0.569(±0.001), 49.9(±0.210) and 50.1(±0.210) respectively. On comparing the mean, CAI, GC and AT of Nucleocapsid of all six strains, the values of optimized DNA was found to be significantly higher. The mean CAI and GC in optimized DNA were found to be 2.3 (135.1%) and 1.2(9.9%) fold higher than respective mean values of wild-type. However, mean of AT content in optimized DNA was decreased by 8.2 % compared to wild-type (**Table 1**). A graph was then plotted taking the CAI values on the x-axis while the number of strains studied on y-axis (**Fig. 1**). The nucleocapsid gene sequences of the wild-type and codon-optimized were aligned as shown (**Fig. S1**). Codon optimization did not change the amino acid sequence of nucleocapsid protein.

**Figure 1:**
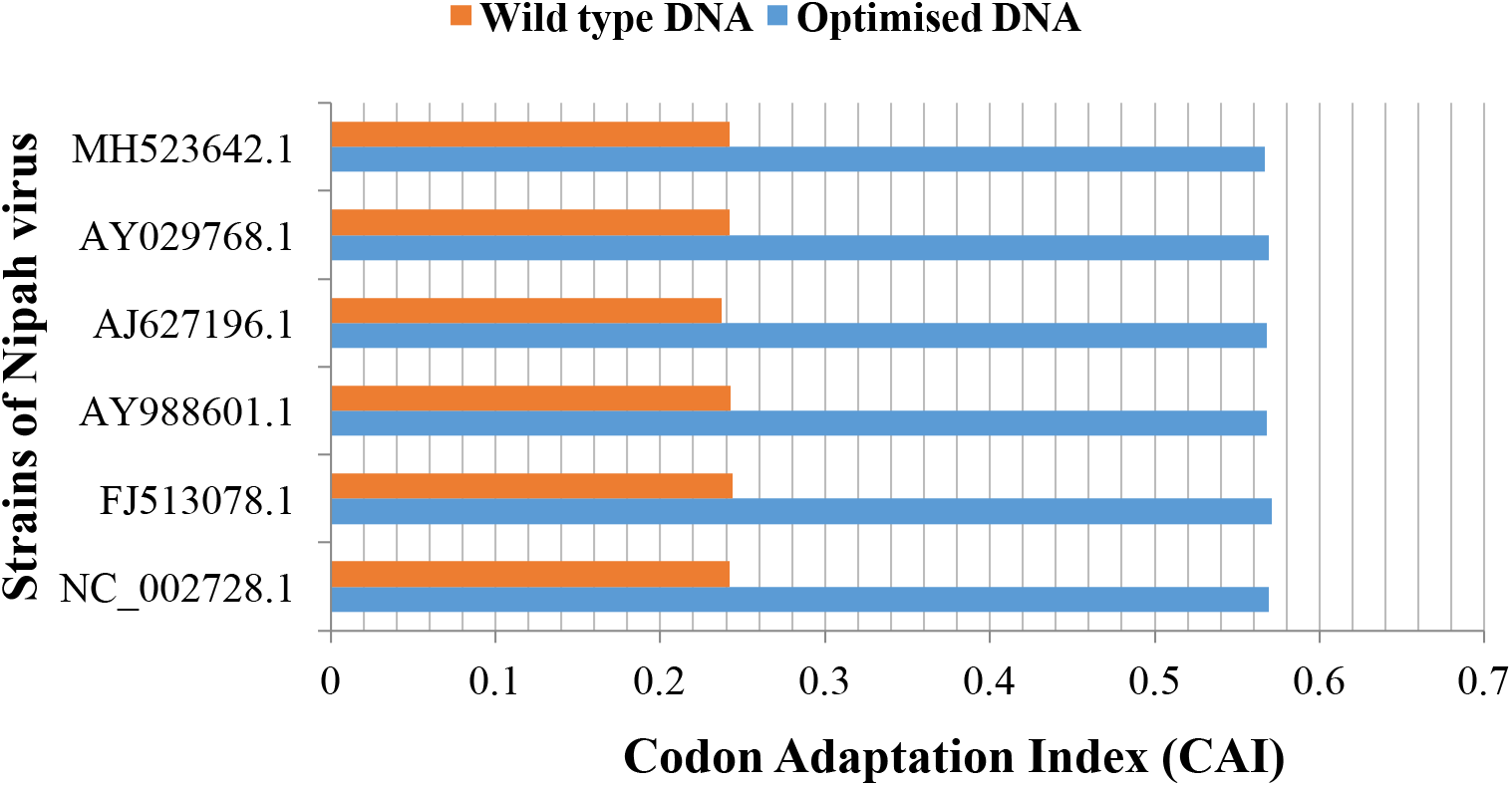
Comparison of the wild-type and optimized DNA sequences of nucleocapsid gene.

Similarly, the CAI, GC and AT frequencies in strains of wild-type P/V/C range from 0.290 to 0.299, 43.3 to 43.7 and 56.3 to 56.7 respectively with an average (±SD) of 0.296(±0.003), 43.4(±0.167) and 56.6(±0.167) respectively. Their respective frequencies in optimized DNA range from 0.577 to 0.595, 46.5 to 47.0 and 53.0 to 53.5 with an average (±SD) of 0.587(±0.007), 46.8(±0.160) and 53.2(±0.160) respectively. The mean CAI and GC in optimized DNA were found to be 2.0 (98.3 %) and 1.1(7.8%) fold higher than respective values of wild-type. Though, mean of AT content in optimized DNA was decreased by 6 % compared to wild-type (**Table 2**). A graph was then plotted for the same (**Fig. 2**). The P/V/C gene sequences of the wild-type and codon-optimized were aligned as presented in (**Fig. S2)**.

**Figure 2:**
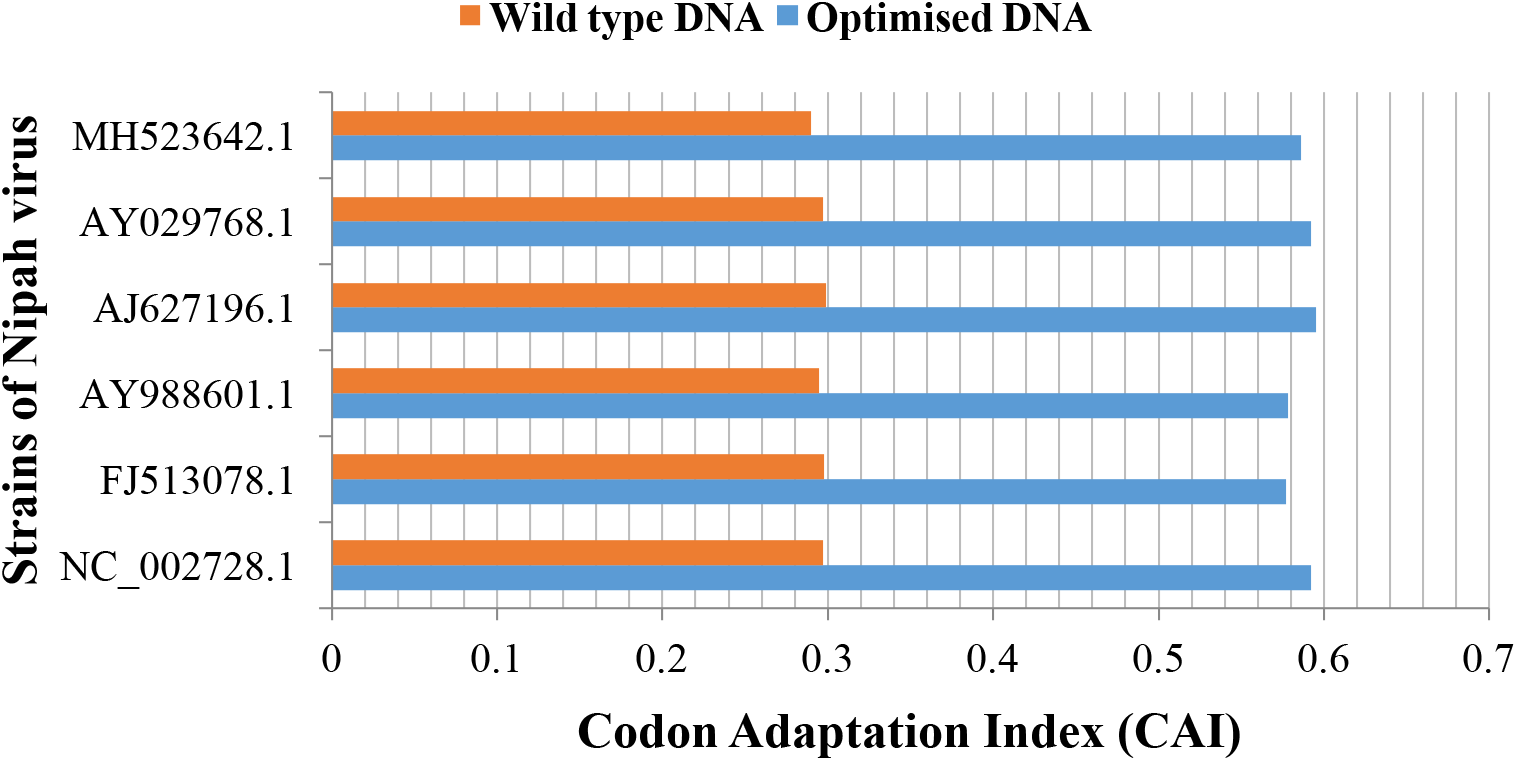
Comparison of the wild-type and optimized DNA sequences of phosphoprotein gene

**TABLE 2:** Expression level of P/V/C (phosphoprotein) gene of Nipah Virus in *E. coli* of wild-type and codon optimized sequences.

| NIPAH VIRUS STRAINS (GenBank Accession No.) | WILD-TYPE DNA |  |  | OPTIMIZED DNA |  |  |
| --- | --- | --- | --- | --- | --- | --- |
|  | CAI | GC % | AT% | CAI | GC % | AT% |
| NC_002728.1 | 0.297 | 43.3 | 56.7 | 0.592 | 46.8 | 53.2 |
| FJ513078.1 | 0.298 | 43.3 | 56.7 | 0.577 | 46.5 | 53.5 |
| AY988601.1 | 0.295 | 43.5 | 56.5 | 0.578 | 46.8 | 53.2 |
| AJ627196.1 | 0.299 | 43.3 | 56.7 | 0.595 | 46.8 | 53.2 |
| AY029768.1 | 0.297 | 43.3 | 56.7 | 0.592 | 46.8 | 53.2 |
| MH523642.1 | 0.290 | 43.7 | 56.3 | 0.586 | 47.0 | 53.0 |
| N | 6 | 6 | 6 | 6 | 6 | 6 |
| Min. | 0.290 | 43.3 | 56.3 | 0.577 | 46.5 | 53.0 |
| Max. | 0.299 | 43.7 | 56.7 | 0.595 | 47.0 | 53.5 |
| Mean±SD | 0.296<br>±0.003 | 43.4<br>±0.167 | 56.6<br>±0.167 | 0.587<br>±0.007** | 46.8<br>±0.160** | 53.2<br>s±0.160* |

\*\*– p <0.002: in comparison with wild type.

Further, the CAI, GC and AT frequencies of matrix protein in wild-type strains range from 0.287 to 0.300, 42.6 to 42.9 and 57.1 to 57.4 respectively with an average (±SD) of 0.294(±0.005), 42.8(±0.138) and 57.3(±0.138) respectively. The respective frequencies of these in optimized DNA range from 0.566 to 0.606,45.5 to 46.1 and 53.9 to 54.5 with an average (±SD) of 0.585(±0.020), 45.9(±0.197) and 54.2(±0.197). The mean CAI and GC in optimized DNA were found to be 2.0(99.0%) and 1.1(7.2%) fold higher than respective wild-type strain. But, mean of AT content in optimized DNA was decreased by 5.4 % compared to wild-type (**Table 3**). A Graph was plotted similar to above mentioned graphs **(Fig. 3)**. The matrix protein gene sequences of the wild-type and codon-optimized were aligned as shown (**Fig. S3**). Codon optimization did not show any modification in the amino acid sequence of matrix protein.

**TABLE 3:** Expression level of M (matrix protein) geneof Nipah Virus in *E. coli* of wild-type and codon-optimized sequences.

| NIPAH VIRUS STRAINS (GenBank Accession No.) | WILD-TYPE DNA |  |  | OPTIMIZED DNA |  |  |
| --- | --- | --- | --- | --- | --- | --- |
|  | CAI | GC % | AT% | CAI | GC % | AT% |
| NC_002728.1 | 0.287 | 42.6 | 57.4 | 0.566 | 45.9 | 54.1 |
| FJ513078.1 | 0.299 | 42.8 | 57.2 | 0.604 | 45.8 | 54.2 |
| AY988601.1 | 0.300 | 42.9 | 57.1 | 0.606 | 45.9 | 54.1 |
| AJ627196.1 | 0.291 | 42.9 | 57.1 | 0.568 | 46.1 | 53.9 |
| AY029768.1 | 0.287 | 42.6 | 57.4 | 0.566 | 45.9 | 54.1 |
| MH523642.1 | 0.297 | 42.7 | 57.3 | 0.601 | 45.5 | 54.5 |
| N | 6 | 6 | 6 | 6 | 6 | 6 |
| Min. | 0.287 | 42.6 | 57.1 | 0.566 | 45.5 | 53.9 |
| Max. | 0.300 | 42.9 | 57.4 | 0.606 | 46.1 | 54.5 |
| Mean±SD | 0.294<br>±0.005 | 42.8<br>±0.138 | 57.3<br>±0.138 | 0.585<br>±0.020** | 45.9<br>±0.197** | 54.2<br>±0.197** |

\*\* – p < 0.002: in comparison with wild type.

**Figure 3:**
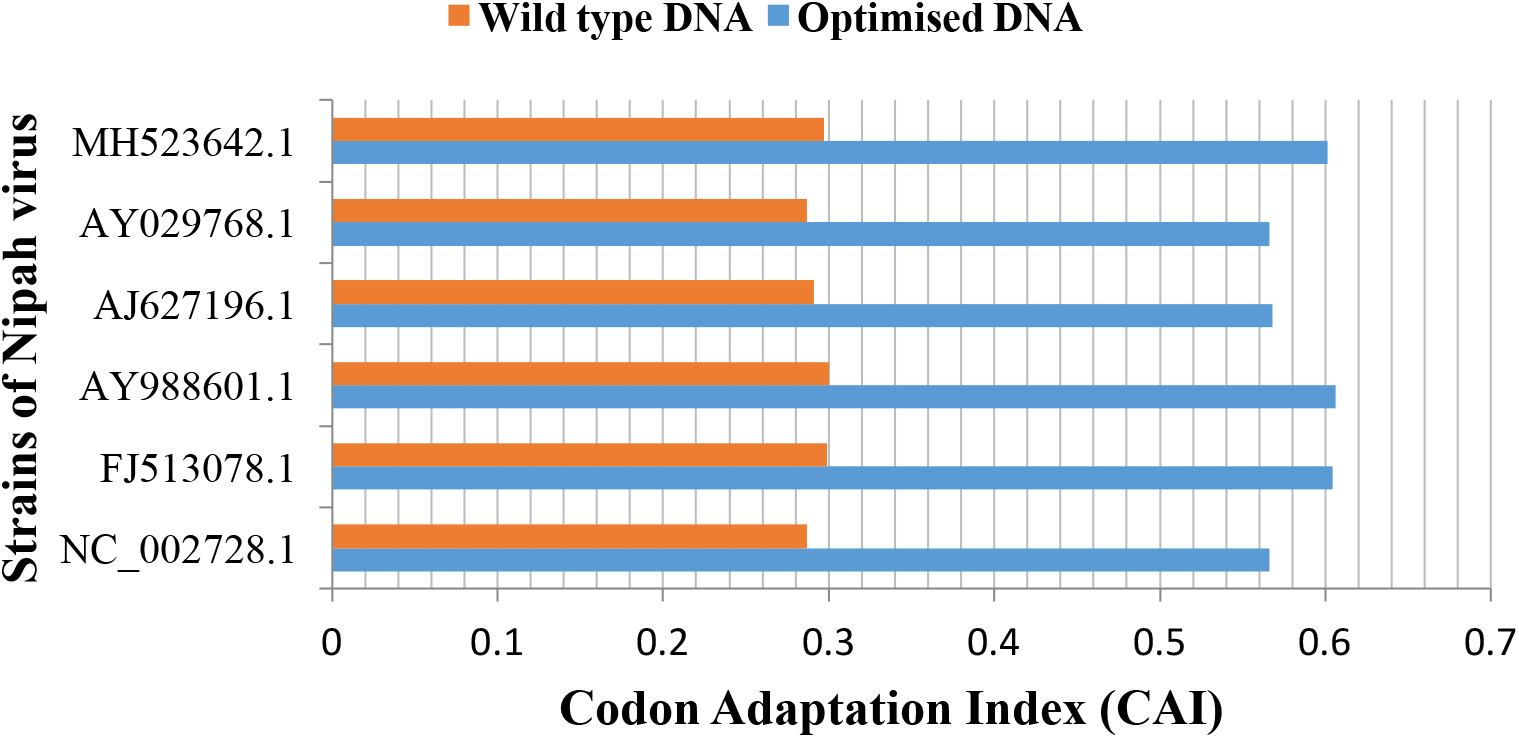
Comparison of the wild-type and optimized DNA sequences of matrix gene.

Furthermore, the CAI, GC and AT frequencies of fusion protein in wild-type strains range from 0.248 to 0.250, 37.9 to 38.5 and 61.5 to 62.1 respectively with an average (±SD) of 0.249(±0.001), 38.2(±0.240) and 61.8(±0.240) respectively; while these values for attachment glycoprotein range from 0.258 to 0.286, 39.8 to 40.4 and 59.6 to 60.2 respectively with an average (±SD) of 0.271(±0.014), 40.0(±0.228) and 60.0(±0.228) respectively. Their respective frequencies in optimized DNA range from 0.597 to 0.607, 43.9 to 44.3 and 55.7 to 56.1 with an average (±SD) of 0.604(±0.003), 44.1(±0.163) and 55.9(±0.163). The mean CAI and GC in optimized DNA were found to be 2.4 (142.5 %) and 1.2 (15.4%) fold higher than respective wild-type strain. However, mean of AT content in optimized DNA was decreased by 9.5 % compared to wild-type (**Table 4**).

**TABLE 4:**
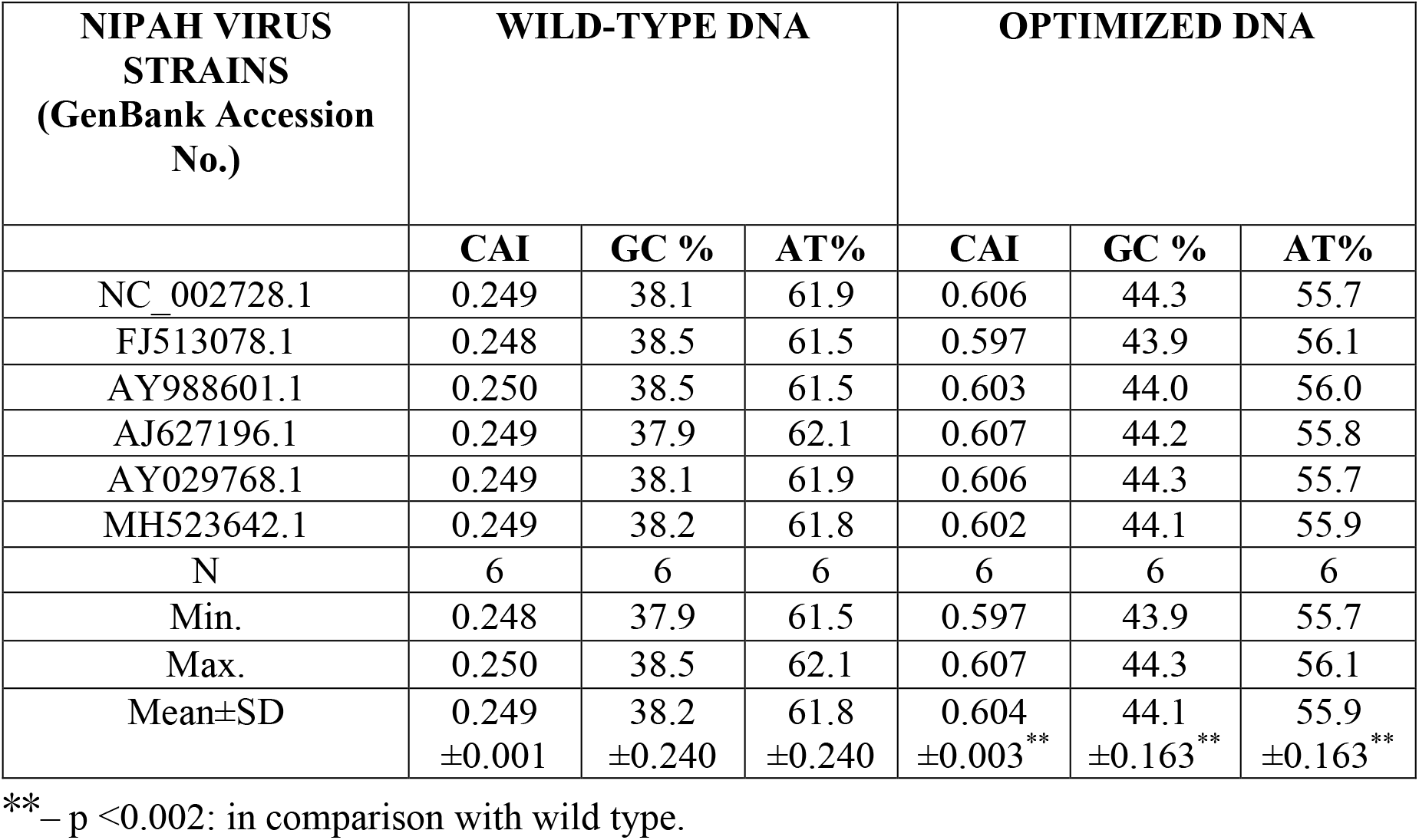
Expression level of F (fusion protein) gene of Nipah Virus in *E. coli* of wild-type and codon optimized sequences.

| NIPAH VIRUS STRAINS (GenBank Accession No.) | WILD-TYPE DNA |  |  | OPTIMIZED DNA |  |  |
| --- | --- | --- | --- | --- | --- | --- |
|  | CAI | GC % | AT% | CAI | GC % | AT% |
| NC_002728.1 | 0.249 | 38.1 | 61.9 | 0.606 | 44.3 | 55.7 |
| FJ513078.1 | 0.248 | 38.5 | 61.5 | 0.597 | 43.9 | 56.1 |
| AY988601.1 | 0.250 | 38.5 | 61.5 | 0.603 | 44.0 | 56.0 |
| AJ627196.1 | 0.249 | 37.9 | 62.1 | 0.607 | 44.2 | 55.8 |
| AY029768.1 | 0.249 | 38.1 | 61.9 | 0.606 | 44.3 | 55.7 |
| MH523642.1 | 0.249 | 38.2 | 61.8 | 0.602 | 44.1 | 55.9 |
| N | 6 | 6 | 6 | 6 | 6 | 6 |
| Min. | 0.248 | 37.9 | 61.5 | 0.597 | 43.9 | 55.7 |
| Max. | 0.250 | 38.5 | 62.1 | 0.607 | 44.3 | 56.1 |
| Mean±SD | 0.249<br>±0.001 | 38.2<br>±0.240 | 61.8<br>±0.240 | 0.604<br>±0.003** | 44.1<br>±0.163** | 55.9<br>±0.163** |

\*\* – p < 0.002: in comparison with wild type.

Whereas the respective frequencies for attachment glycoprotein in optimized DNA range from 0.560 to 0.602, 44.2 to 44.9 and 55.1 to 55.8 with an average (±SD) of 0.582(±0.020), 44.5(±0.308) and 55.5(±0.308) respectively. The mean CAI and GC in optimized DNA were found to be 2.1(114.8 %) and 1.1(11.2%) fold higher than respective wild-type values. However, mean of AT content in optimized DNA was decreased by 7.5 % compared to wild-type (**Table 5**). The graphs were then plotted for both the genes taking the CAI values on the x-axis and the number of strains studied on y-axis **(Fig. 4 and 5)**. The fusion protein gene sequences of the wild-type and codon-optimized were aligned as shown in (**Fig. S4**). The glycoprotein gene sequences of the wild-type and codon-optimized were aligned as presented (**Fig. S5**). Codon optimization did not alter the amino acid sequence of glycoprotein.

**Figure 4:**
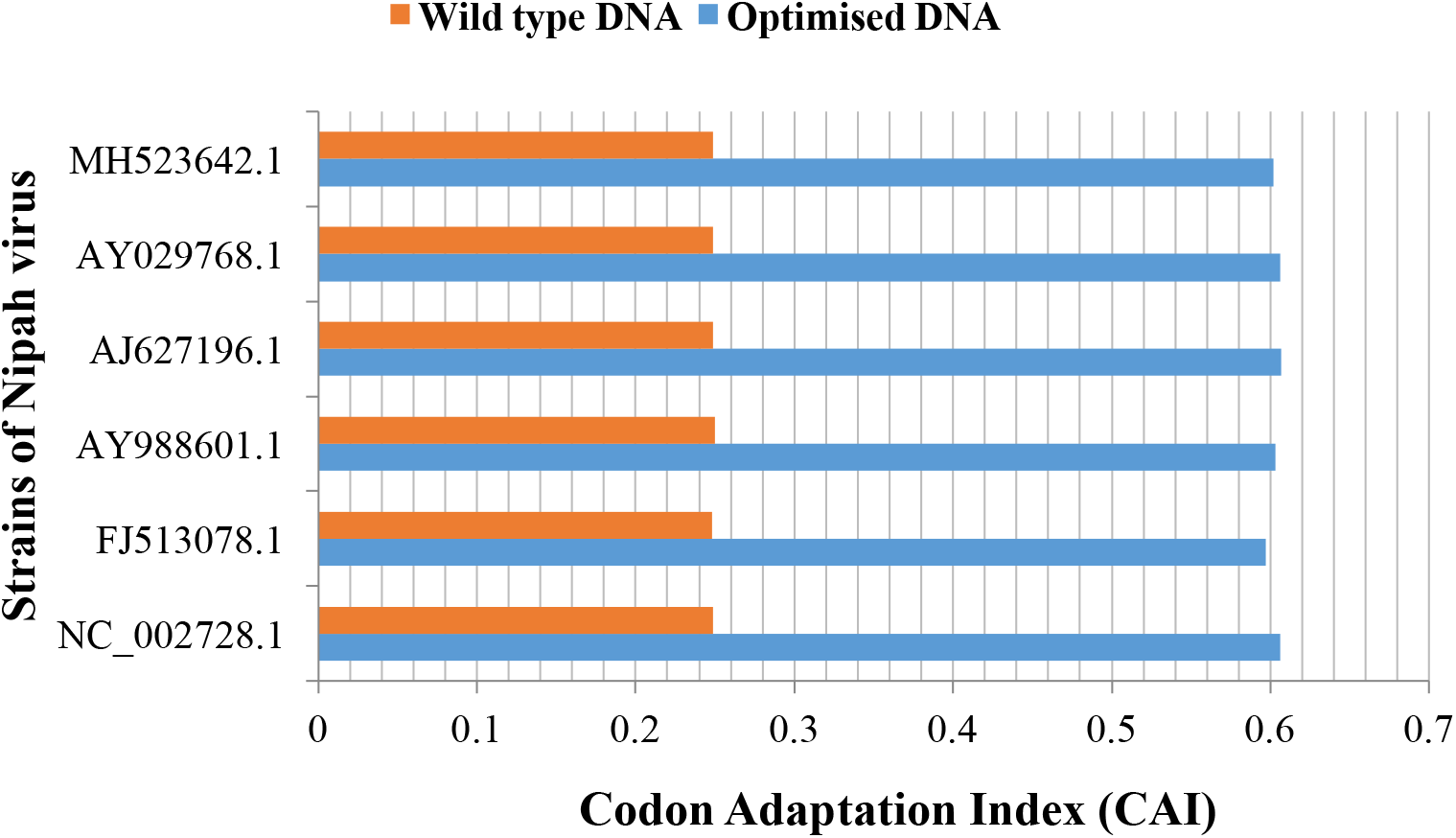
Comparison of the wild-type and optimized DNA sequences of fusion protein gene.

**Figure 5:**
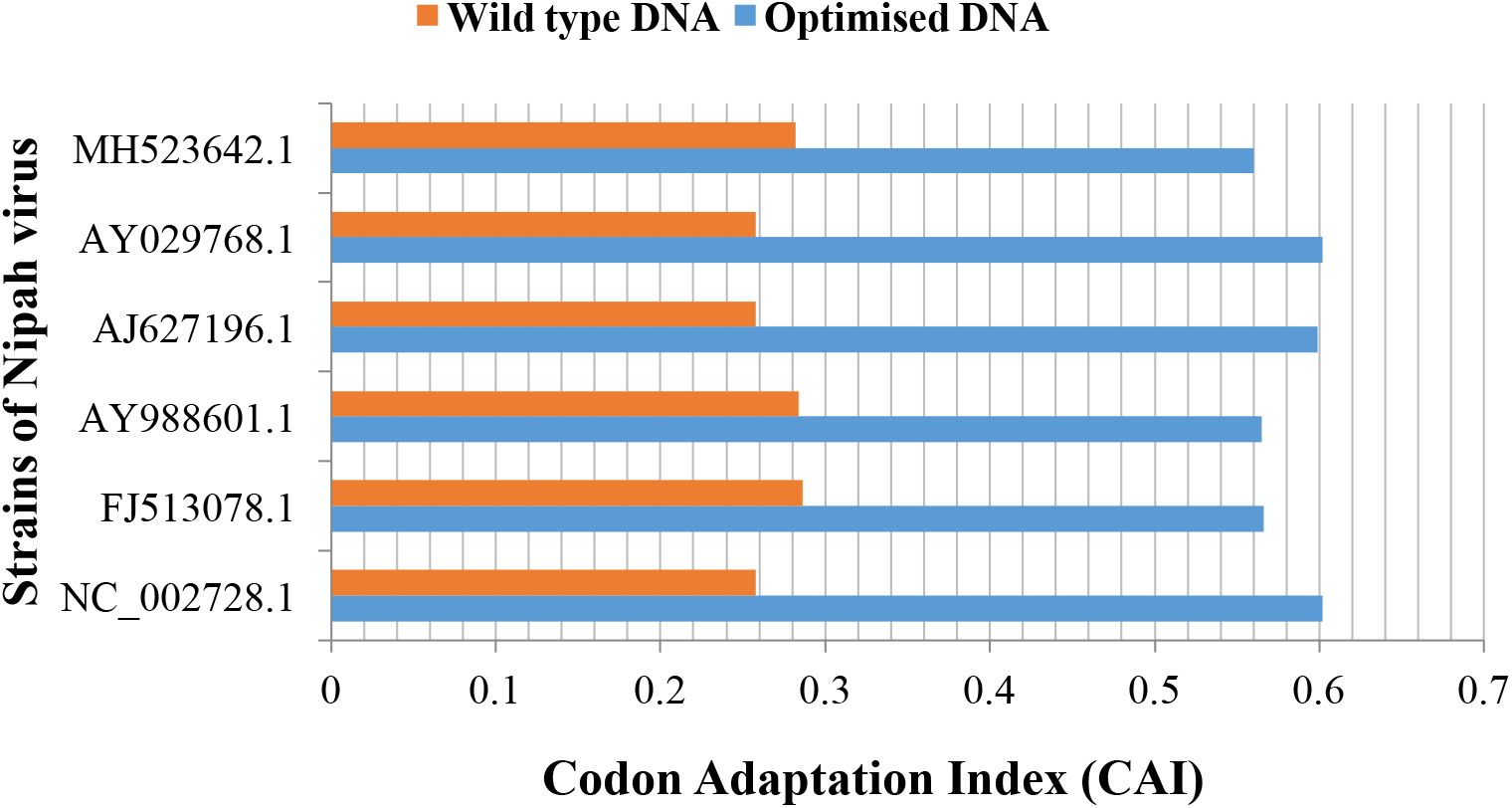
Comparison of the wild-type and optimized DNA sequences of attachment glycoprotein.

**TABLE 5:** Expression level of G (attachment glycoprotein) gene of Nipah Virus in *E. coli* of wild-type and codon optimized sequences.

| NIPAH VIRUS STRAINS<br>(GenBank Accession No.) | WILD-TYPE DNA |  |  | OPTIMIZED DNA |  |  |
| --- | --- | --- | --- | --- | --- | --- |
|  | CAI | GC % | AT% | CAI | GC % | AT% |
| NC_002728.1 | 0.258 | 39.8 | 60.2 | 0.602 | 44.2 | 55.8 |
| FJ513078.1 | 0.286 | 40.1 | 59.9 | 0.566 | 44.7 | 55.3 |
| AY988601.1 | 0.284 | 40.0 | 60.0 | 0.565 | 44.6 | 55.4 |
| AJ627196.1 | 0.258 | 39.9 | 60.1 | 0.599 | 44.2 | 55.8 |
| AY029768.1 | 0.258 | 39.8 | 60.2 | 0.602 | 44.2 | 55.8 |
| MH523642.1 | 0.282 | 40.4 | 59.6 | 0.560 | 44.9 | 55.1 |
| N | 6 | 6 | 6 | 6 | 6 | 6 |
| Min. | 0.258 | 39.8 | 59.6 | 0.560 | 44.2 | 55.1 |
| Max. | 0.286 | 40.4 | 60.2 | 0.602 | 44.9 | 55.8 |
| Mean±SD | 0.271<br>±0.014 | 40.0<br>±0.228 | 60.0<br>±0.228 | 0.582<br>±0.020** | 44.5<br>±0.308** | 55.5<br>±0.308** |

\*\*– p < 0.002: in comparison with wild type.

The CAI, GC and AT frequencies of polymerase (gene L) in wild type DNA range from 0.234 to 0.242,37.5 to 38.1 and 61.9 to 62.5 respectively and an average (±SD) of 0.238(±0.003), 37.8(±0.297), 62.2(±0.297) respectively. The respective frequencies of these in optimized DNA range from 0.579 to 0.581, 44.1 to 44.4 and 55.6 to 55.9 with an average (±SD) of 0.580(±0.001), 44.3(±0.164), 55.8(±0.164). The mean CAI and GC in optimized DNA were found to be 2.4(143.7%) and 1.2(17.2%) fold higher than respective wild-type values. However, mean of AT content in optimized DNA was reduced by 10.3 % compared to wild-type (**Table 6**). A graph was plotted on similar basis for all strains **(Fig. 6)**. The polymerase gene sequences of the wild-type and codon-optimized were aligned as shown (**Fig. S6**). It aims to increase the immunogenicity of epitope-based vaccines as it can enhance translational efficiency. Thus, modification of the codon bias of gene sequences is a promising tool of gene expression control.

**TABLE 6:** Expression level of L (polymerase) gene of Nipah Virus in *E. coli* of wild-type and codon optimized sequences.

| NIPAH VIRUS STRAINS<br>(GenBank Accession No.) | WILD-TYPE DNA |  |  | OPTIMIZED DNA |  |  |
| --- | --- | --- | --- | --- | --- | --- |
|  | CAI | GC % | AT% | CAI | GC % | AT% |
| NC_002728.1 | 0.234 | 37.6 | 62.4 | 0.581 | 44.1 | 55.9 |
| FJ513078.1 | 0.24 | 38 | 62 | 0.58 | 44.4 | 55.6 |
| AY988601.1 | 0.241 | 38.1 | 61.9 | 0.579 | 44.4 | 55.6 |
| AJ627196.1 | 0.234 | 37.5 | 62.5 | 0.581 | 44.1 | 55.9 |
| AY029768.1 | 0.234 | 37.5 | 62.5 | 0.581 | 44.1 | 55.9 |
| MH523642.1 | 0.242 | 38.1 | 61.9 | 0.579 | 44.4 | 55.6 |
| N | 6 | 6 | 6 | 6 | 6 | 6 |
| Min. | 0.234 | 37.5 | 61.9 | 0.579 | 44.1 | 55.6 |
| Max. | 0.242 | 38.1 | 62.5 | 0.581 | 44.4 | 55.9 |
| Mean±SD | 0.238<br>±0.003 | 37.8<br>±0.297 | 62.2<br>±0.297 | 0.580<br>±0.001** | 44.3<br>±0.164** | 55.8<br>±0.164** |

\*\*– p < 0.002: in comparison with wild type.

**Figure 6:**
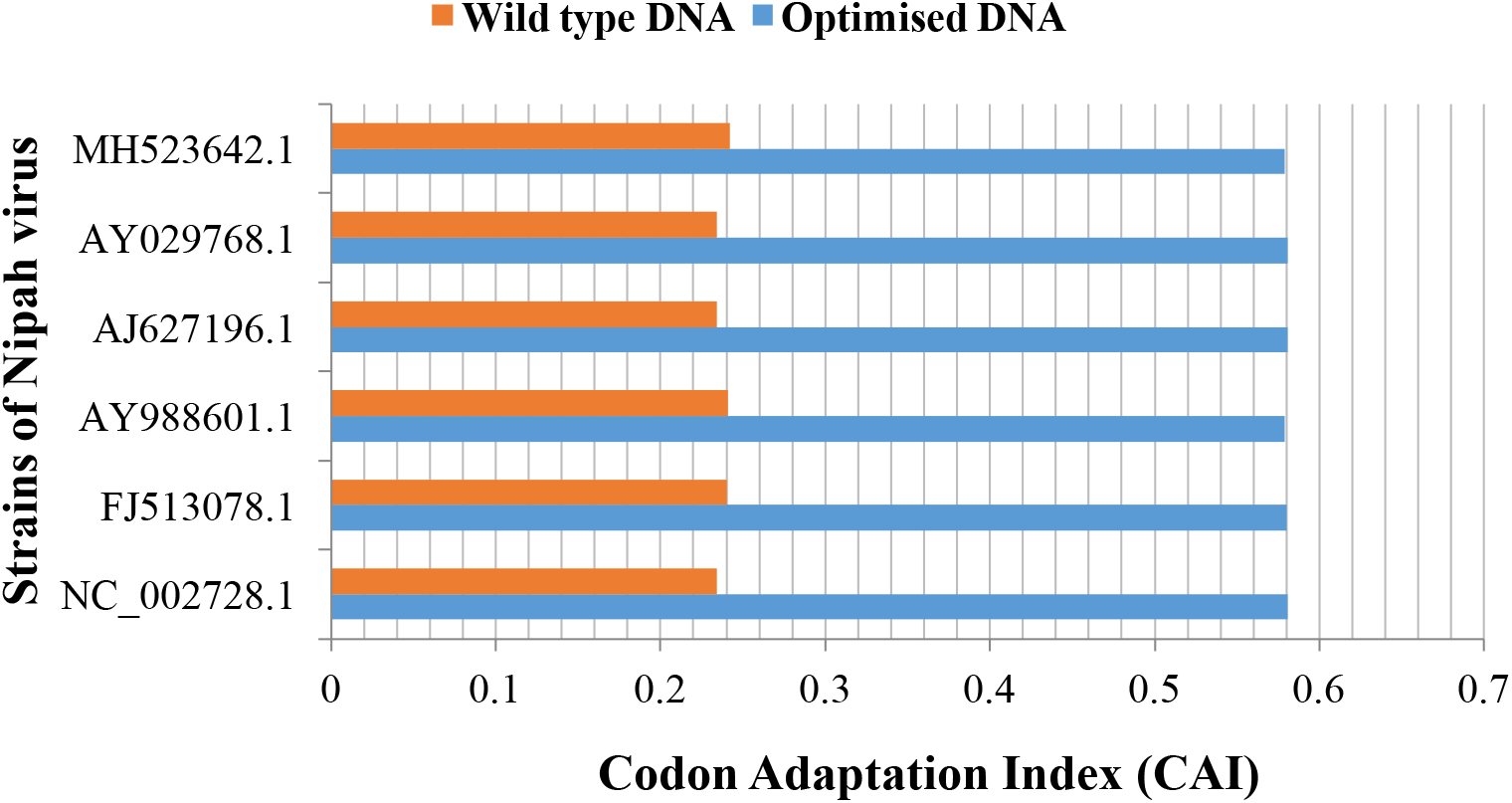
Comparison of the wild-type and optimized DNA sequences of polymerase gene.

## Discussion

The applications of DNA codon optimization range from numerous animal tests to remove stop codons, to clone, in custom design of synthetic genes, to improve the functionality of genes, to increase protein expression level, for lower production costs, as well as in drug development. The promise of DNA based immunity has been indicated by few human trials for HIV infection. Codon usage adaptation of the gag protein of HIV delivered by a DNA vaccine increased gene expression by 10-fold compared to wild-type [26]. Further, gene optimization has been effective for a number of treatment applications where a protein is synthesized *in vivo* following gene delivery and is becoming routinely used for a range of applications [27]. For example, codon optimization of the gene for F protein, expressed from a DNA vaccine of the Respiratory syncytial virus improved the performance relative to wild-type [28]. Codon optimization also ensures that the 5’ mRNA end is unlikely to form stable hairpins, thus facilitating optimal mRNA loading and protein translation. This was elegantly shown by expressing 154 green fluorescent protein (GFP) mutants in *E. coli*, where hairpins engineered into the 5’ mRNA end reduced GFP expression by up to 250-fold, compared to an optimal codon-optimized construct [29]. Apart from increasing protein expression levels, codon usage also finds application in metagenomic studies. RNA virus genomes from high temperature acidic metagenomes have been analysed on the basis of codon usage frequency to determine their host range, whether bacterial, archaeal or eukaryal [30].

In the present study, the mean, CAI and GC of all NiV strains, the values of optimized DNA were found to be significantly different and higher than their respective wild-type strain, in case of all genes. On an average, CAI of N gene in optimized DNA was enhanced by 2.3 (135.1%) fold, while in P/V/C, it was increased by 2.0 (98.3%) fold, respectively. Further, the CAI in optimized DNA of M gene and F gene was enhanced by 2.0 (99.0%) fold for gene M and 2.4 (142.5 %), fold for gene F. Gene G showed an increase of 2.1 (114.8 %) fold, and gene L showed an increase of 2.4 (143.7%) fold. Also, an increase in the percentage of GC content was observed in optimized DNA sequences as compared to the wild-type sequences. Further, on an average GC content of the N gene in optimized DNA was enhanced by 1.2(9.9 %) fold, while in P/V/C, it was increased by 1.1 (7.8%) fold, respectively. Similarly, GC content in an optimized DNA of M gene and F gene was increased by 2.4 (142.5 %) and 1.2 (15.4%) fold, respectively. Gene G showed an increase of 1.1(11.2%) fold for GC content, and gene L showed an increase of 1.2 (17.2%) fold. However, AT content in all genes was significantly decreased, as compare to wild-type.

A handful of vaccine candidates are in development that employ NiV glycoprotein (G) and fusion (F) proteins to stimulate a protective immune response in preclinical animal models. Some approaches target specific neutralizing antibody responses; others have been evaluated for both immune response and efficacy [31]. NiV envelope glycoprotein G was found to effectively induce specific antibody responses which could block NiV entry to susceptible cells [32]. The generation of neutralizing antibodies in response to G protein is suggestive of a robust adaptive immune response, which is an essential prerequisite of a good vaccine. While G may itself serve as a good vaccine antigen, the stable prefusion of both F and G antigens has shown to produce a higher and broader multivalent polyclonal antibody response in mice models, making it a more potent candidate for vaccine development [33]. Therefore, G protein or G/F prefusions, being more immunogenic, can serve as good vaccine candidates. Further, codon optimization can be used to increase the expression of these genes in cells in order to achieve high titres for large scale production. Thus, codon optimization coupled with immunology-based studies is important to produce effective vaccines, which have a potential of being upscaled at industrial level. Identifying and producing such vaccines will provide an excellent therapeutic strategy for fighting Nipah virus infection.

Nonetheless, codon optimization might pose certain challenges. Although codon optimization has applications like recombinant protein drugs and nucleic acid therapies, including gene therapy, mRNA therapy, and DNA/RNA vaccine, recent reports indicate that it can affect protein conformation and function, reduce efficacy and increase immunogenicity. It may decrease the safety and efficacy of biotech therapeutics [34]. Synonymous codon changes may affect protein conformation and stability, change sites of post translational modifications, and alter protein function. Moreover, synonymous mutations have been linked to numerous diseases. The effects of synonymous codon changes were highlighted in a recent study where the fluorescent properties of a protein were altered by synonymous codon changes due to altered protein folding [35]. Thus, *in vitro* analysis of such in silico studies is required to overcome these challenges.

## Conclusion

A range of clinical presentations result from Nipah virus infections in humans, from asymptomatic infection (subclinical) to acute respiratory infection and fatal encephalitis. The case fatality rate is roughly calculated at 40% to 75%. There is no treatment or vaccine available for either people or animals [24]. High pathogenicity of Nipah virus in humans and lack of appropriate immunological based therapeutics and diagnosis for prevention and cure of the disease, accounts for the need of investigators worldwide to develop efficient vaccine and treatment regimes. Vaccines are generally proteins with immunogenic properties and are not expressed in sufficient quantities because of codon bias in the expression host. Thus, the study was carried out to optimize the codon for overexpression of different Nipah virus genes in *E. coli* which could be used to develop vaccine and immunoassay based diagnostic kit. The CAI and GC content of optimized sequences were increased as compared to the wild-type sequences indicating that they can be over-expressed in *E. coli*. Based on our codon optimization study and previous studies on immunogenicity of the proposed genes, we believe that G proteins or F/G fusions can potentially serve as ideal candidates for Nipah Virus vaccine. Future work involves *in-vitro* validation of this in silico study to determine the level of overexpression as well as testing, their safety, and potency in generating an immune response, which can then be applied on an industrial scale for development of immunodiagnostics and immunotherapeutics.

## Acknowledgments

Authors are thankful to the Principal, Gargi College for providing the infrastructural support.

## Legends to supplementary (S) figures

**Figure S1:**
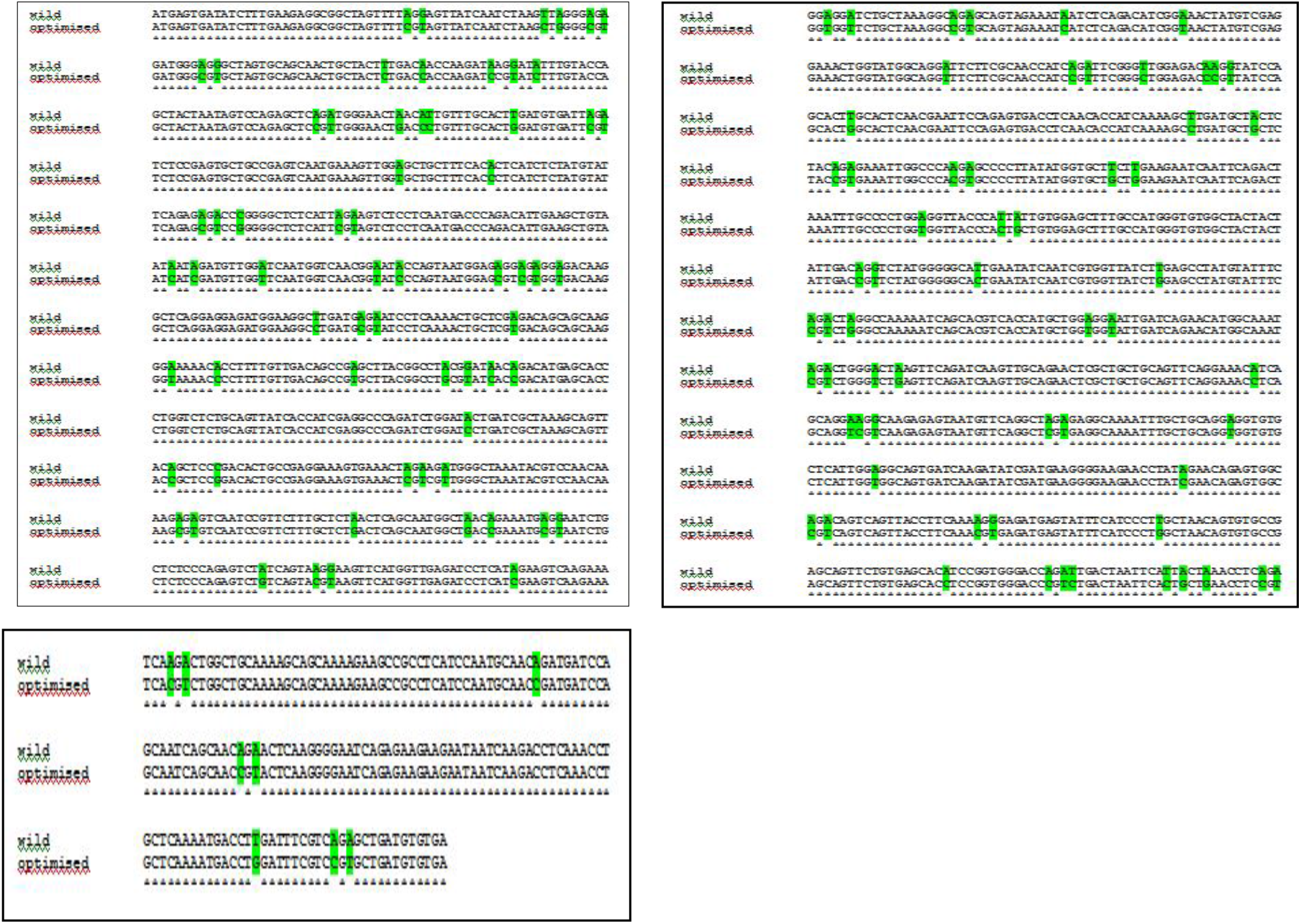
Alignment of wild-type and codon-optimized DNA of nucleocapsid gene of Nipah virus (NC_002728.1). *The bases highlighted in green represent differences in base pairs among the wild-type and optimized sequences.

**Figure S2:**
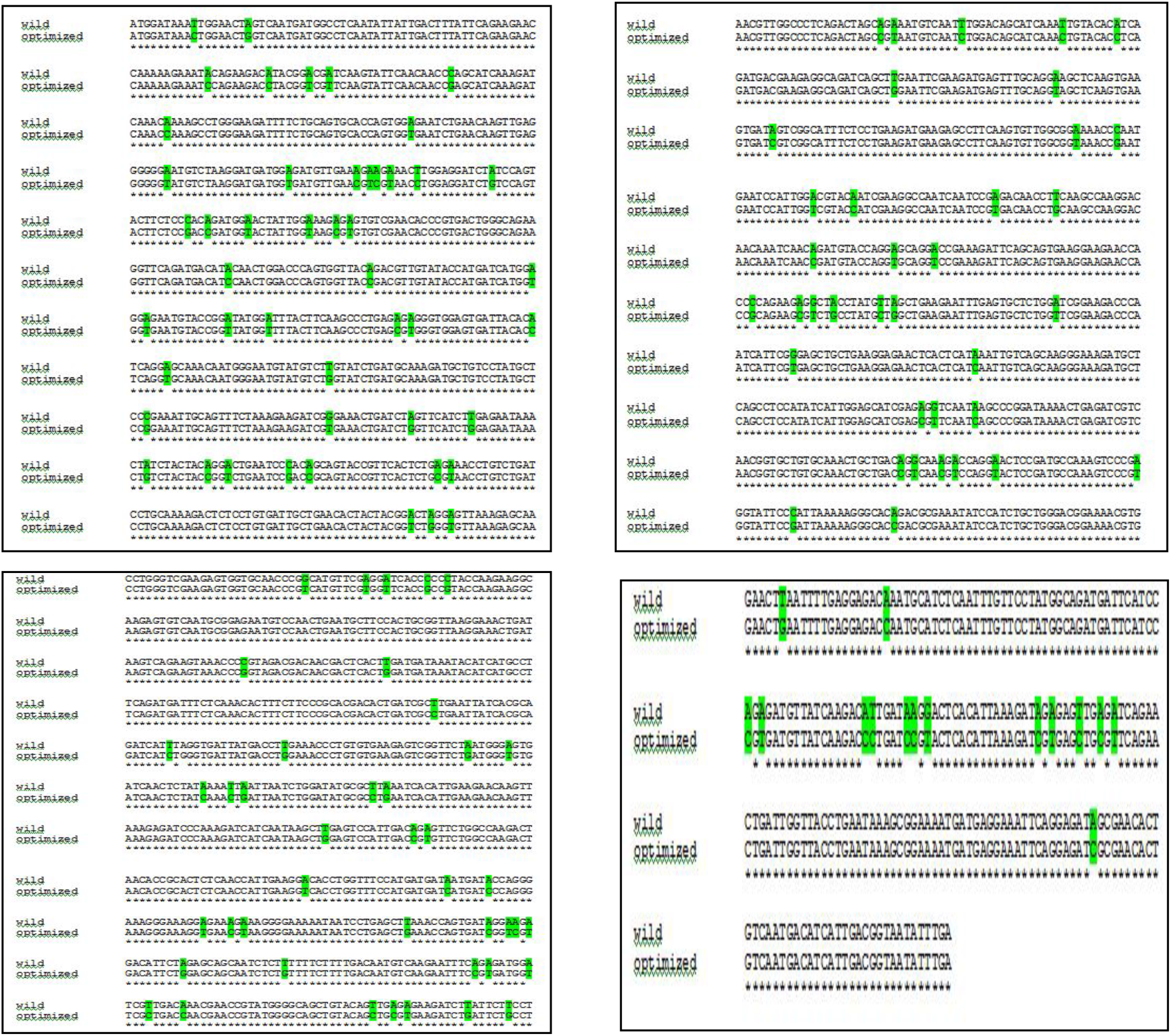
Alignment of wild-type and codon-optimized DNA of P/V/C gene of Nipah virus (NC_002728.1). *The bases highlighted in green represent differences in base pairs among the wild-type and optimized sequences.

**Figure S3:**
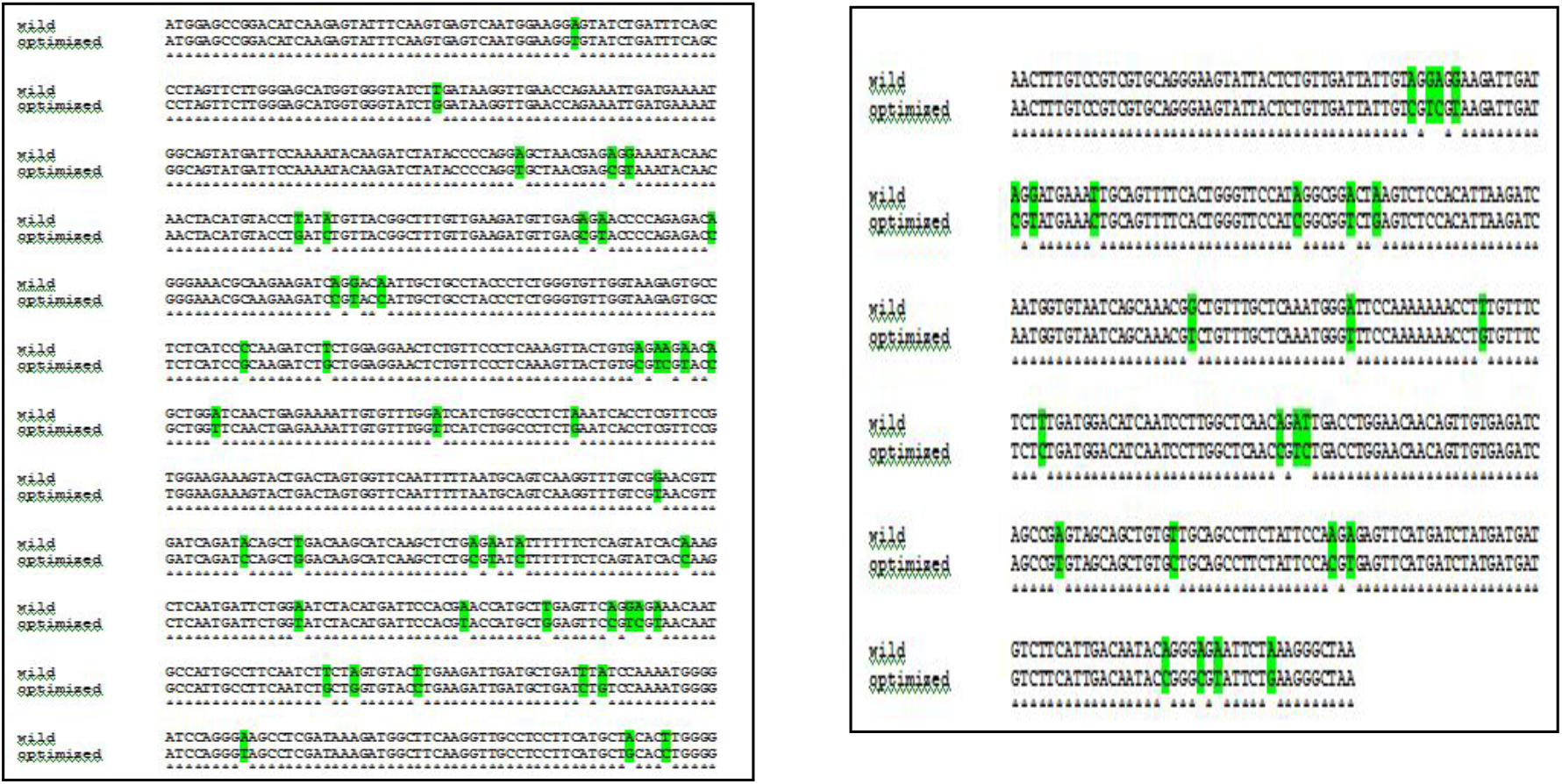
Alignment of wild-type and codon-optimized DNA of matrix protein gene of Nipah virus(NC_002728.1). *The bases highlighted in green represent differences in base pairs among the wild-type and optimized sequences.

**Figure S4:**
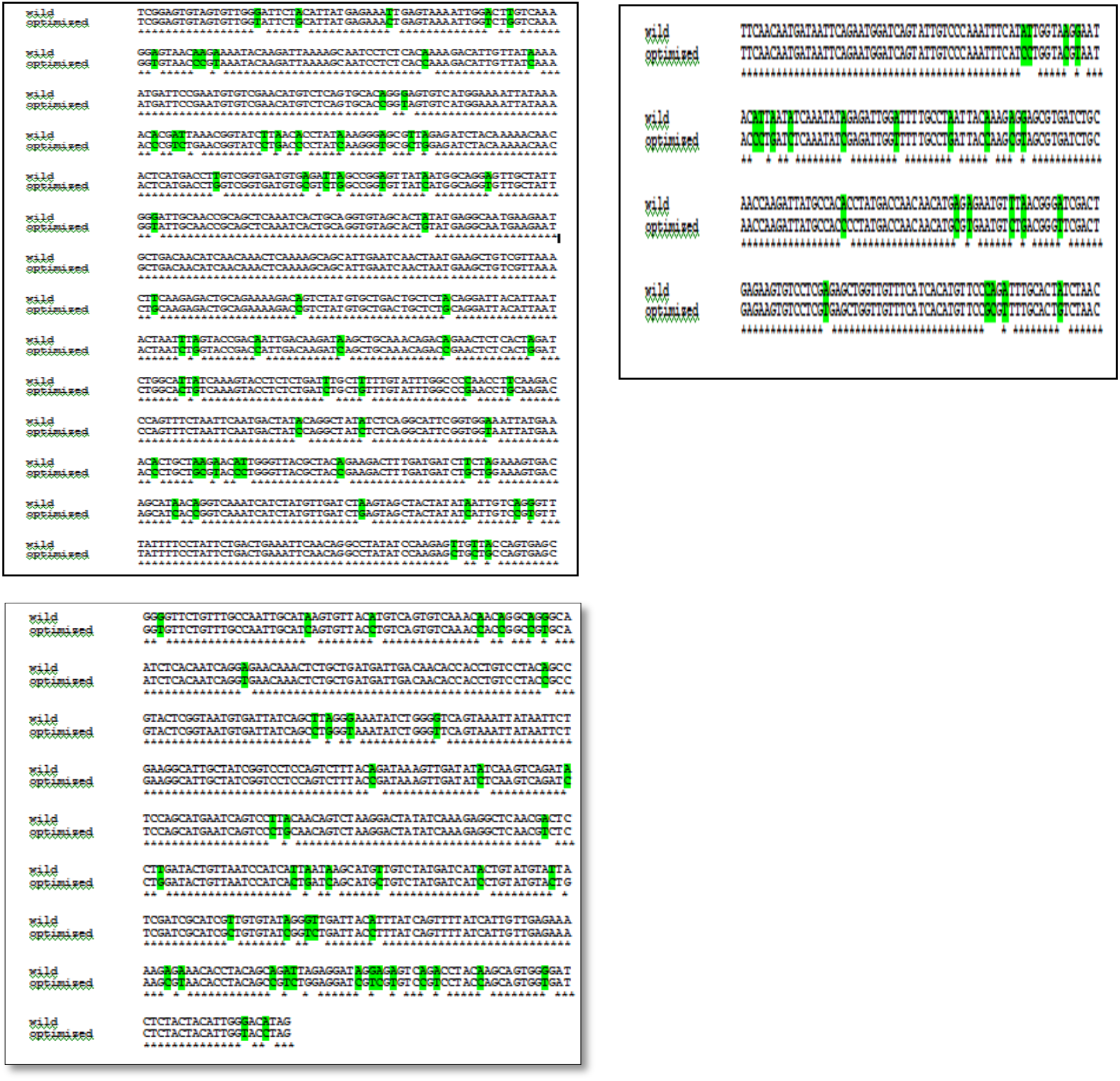
Alignment of wild-type and codon-optimized DNA of fusion protein gene of Nipah virus(NC_002728.1). *The bases highlighted in green represent differences in base pairs among the wild-type and optimized sequences.

**Figure S5:**
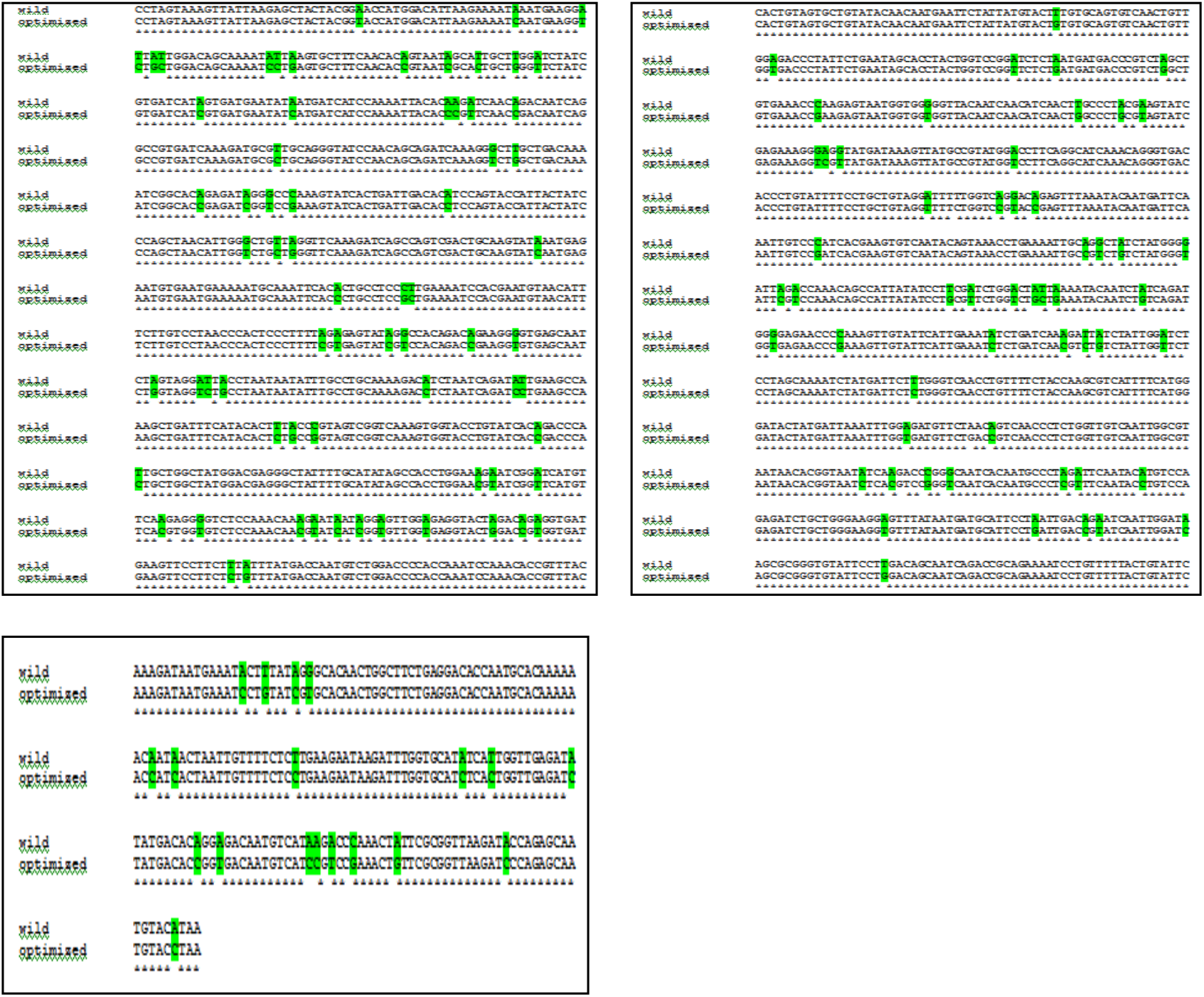
Alignment of wild-type and codon-optimized DNA of glycoprotein gene of Nipah virus. (NC_002728.1). *The bases highlighted in green represent differences in base pairs among the wild-type and optimized sequences.

**Figure S6:**
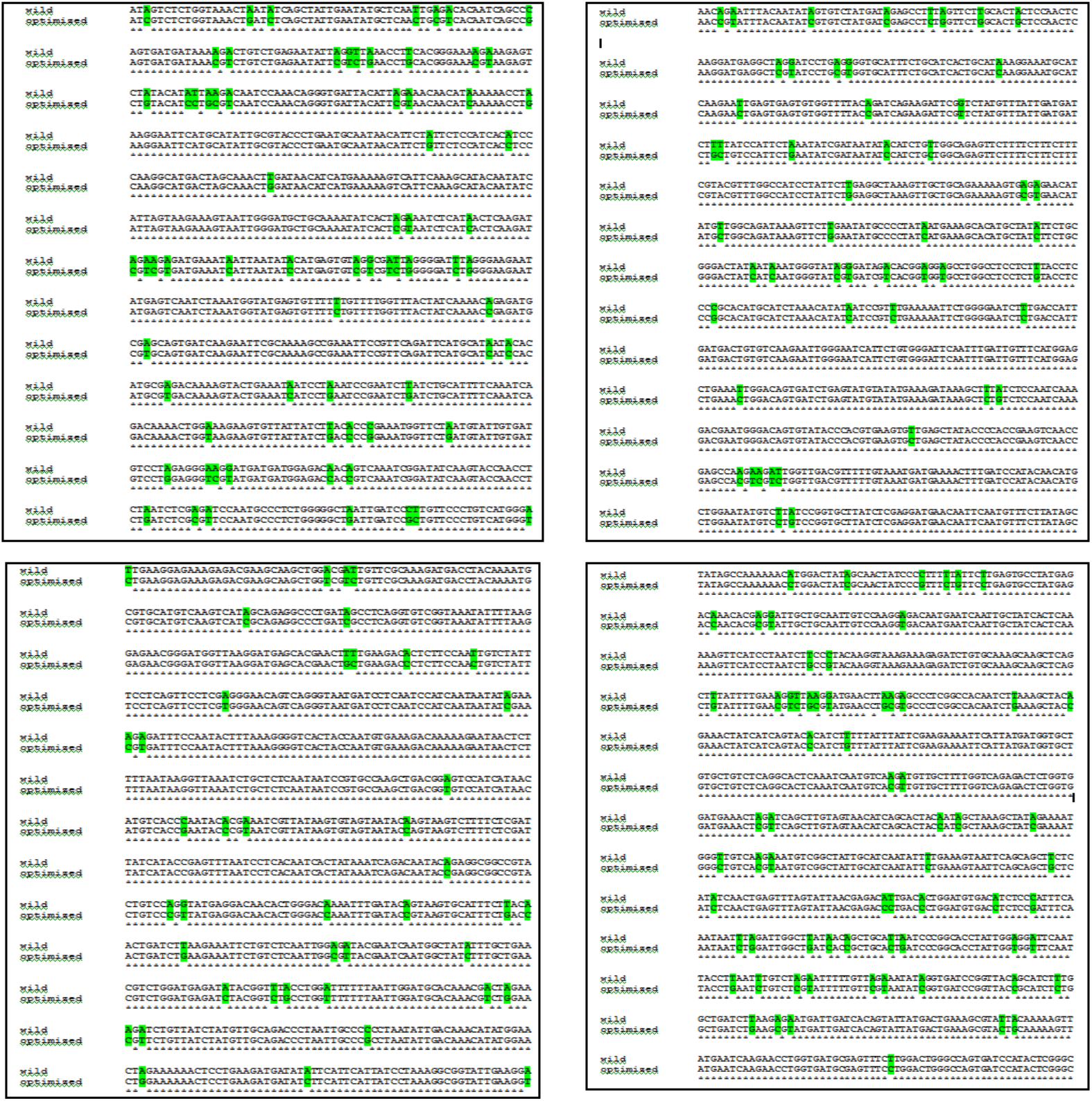
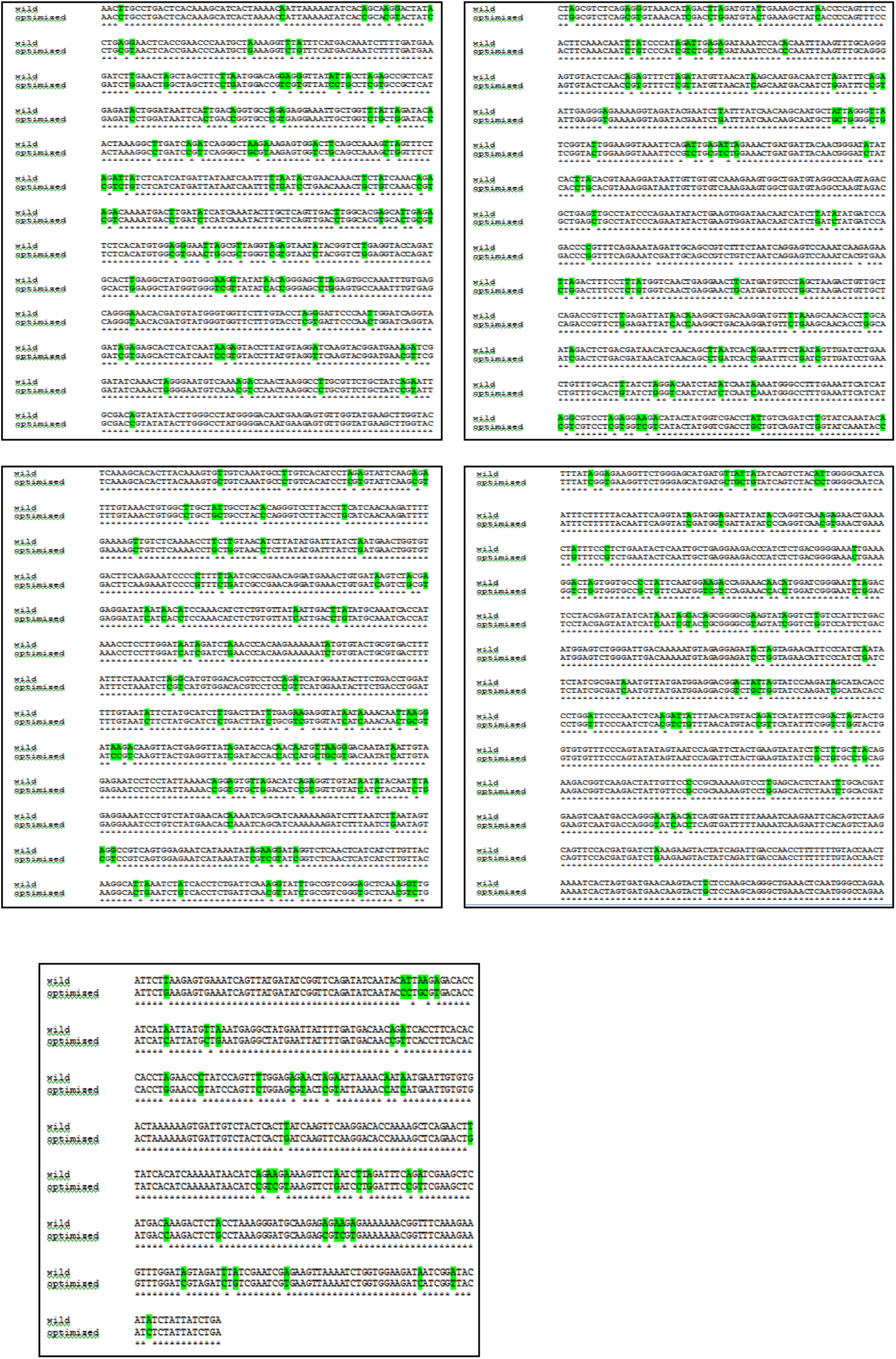
Alignment of wild-type and codon-optimized DNA of polymerase gene of Nipah virus. (NC_002728.1). *The bases highlighted in green represent differences in base pairs among the wild-type and optimized sequences.

